# Distinct developmental organization of intrinsic brain dynamics supports different components of inhibitory control during adolescence

**DOI:** 10.64898/2026.11.17.767444

**Authors:** Da Chang, Donghui Song, Henyi Rao

**Affiliations:** College of Education Science, Hengyang Normal University, Hengyang, China; Department of Psychology, University of Science and Technology of China, Hefei, China

**Keywords:** adolescence, inhibitory control, resting-state functional magnetic resonance imaging, brain entropy, cortical hierarchy, contextual control

## Abstract

Adolescent inhibitory control develops within a changing organization of intrinsic brain activity, yet the signal properties and cortical systems associated with different control demands remain unclear. We studied 101 youths aged 8–18 years using brain entropy (BEN), amplitude of low-frequency fluctuations (ALFF), fractional ALFF (fALFF), and regional homogeneity (ReHo) derived from resting-state functional magnetic resonance imaging. Voxelwise and seven-network models evaluated age associations and task-by-age interactions while adjusting for sex, head motion, and gray-matter anatomy. Spatially standardized ALFF showed age associations in somatomotor and salience/ventral-attention networks. Contextual control, assessed with the AX Continuous Performance Task (AX-CPT), showed the broadest age-dependent coupling: all four spatially standardized measures yielded voxelwise interactions surviving permutation-based cluster false discovery rate correction, and the frontoparietal BEN interaction survived network correction. Better action cancellation was associated with higher frontoparietal BEN without a corrected network age interaction. Independent cortical references further differentiated these patterns: raw BEN age-effect evidence was stronger toward association cortex, whereas the ALFF–AX-CPT age interaction was stronger toward sensorimotor cortex. Additional task interactions aligned with the second and third connectivity gradients. Core voxelwise AX-CPT findings persisted after age trimming. These findings link the differentiation of inhibitory-control demands to the developmental organization of complementary intrinsic signal properties across cortical systems, highlighting contextual control as a prominent locus of age-dependent brain–behavior coupling.

**Highlights:**

- Age shaped links between intrinsic brain activity and contextual control
- Contextual control showed age interactions across four resting-state measures
- Frontoparietal brain entropy related to contextual control and action cancellation
- Selected age and control effects followed distinct cortical organizing axes
- Core contextual-control interactions persisted after excluding age extremes

## 1. Introduction

The development of inhibitory control enables young people to maintain goals, resist distraction, and regulate actions as environmental demands change. Large-scale behavioral evidence identifies a nonlinear course of executive-function maturation, with substantial gains during late childhood and mid-adolescence and more gradual stabilization toward adulthood (Tervo-Clemmens et al., 2023). Within this shared progression, maintaining contextual information, suppressing a reflexive eye movement, cancelling an initiated action, and resolving competing response tendencies place different demands on cognitive systems. Understanding inhibitory-control development therefore requires examining how these related control demands are supported by the developing brain (Luna et al., 2015; Constantinidis and Luna, 2019; Ordaz et al., 2013).

Cognitive development unfolds within a changing hierarchy of cortical systems. In adults, the principal functional-connectivity gradient extends from primary sensory and motor regions to transmodal association cortex (Margulies et al., 2016). Dong et al. (2021) showed that the dominant cortical gradient shifts from a configuration distinguishing sensory and motor systems in childhood toward this adult-like organization during adolescence. The broader sensorimotor–association (S–A) axis integrates several anatomical and functional features into a common reference for cortical hierarchy (Sydnor et al., 2021). Network studies further link developing integration to inhibitory control (Marek et al., 2015), motivating accounts in which improving executive function depends on the maturation of specialization and communication across hierarchically organized systems (Keller et al., 2023).

These findings establish a spatial framework for development, but leave open which local properties of spontaneous activity relate to specific control demands within that framework. Brain entropy (BEN), estimated here using sample entropy, characterizes the temporal irregularity of blood oxygenation level-dependent (BOLD) signals (Wang et al., 2014; Wang, 2021). Amplitude of low-frequency fluctuations (ALFF), fractional ALFF (fALFF), and regional homogeneity (ReHo) characterize fluctuation amplitude, relatively low-frequency spectral content, and local temporal concordance, respectively (Zang et al., 2004, 2007; Zou et al., 2008). Evidence supports examining these properties together. BEN and fALFF show regionally varying positive and negative associations (Song et al., 2019). In a randomized oxytocin study, left temporoparietal BEN increased in younger adults and decreased in older adults, whereas fALFF increased in the same region, particularly in younger adults (Song and Wang, 2026). These distinct responses support the complementary sensitivity of temporal irregularity and relative spectral content to age-dependent functional modulation. They motivate asking whether different signal properties also capture different age-related associations with cognitive control during youth.

Developmental BEN findings further suggest that regional organization deserves attention alongside whole-brain averages. A naturalistic movie-viewing preprint in children aged 3.5–12 years reported regional BEN decreases with age in areas discussed in relation to the action-mode network, despite no significant within-childhood association between age and global mean BEN (Song, 2025). This observation suggests that developmental variation in temporal irregularity can be spatially selective during engagement with a continuous stimulus. A question for resting-state research is whether such regional differentiation also has behavioral relevance: does age shape the association between intrinsic signal properties and particular control demands, and are these associations organized along cortical hierarchy?

Two distinctions are especially relevant to this question. Age-related variation in the level of a brain measure and age-related variation in its association with behavior address different aspects of development. A measure can carry different behavioral relevance across ages even when its network average shows little age variation. In addition, raw and within-brain spatially standardized measures characterize complementary levels of expression: the original metric scale and the prominence of each location relative to the rest of an individual’s brain. Both occur against substantial anatomical maturation. Lifespan brain charts demonstrate nonlinear trajectories of gray-matter volume (Bethlehem et al., 2022), motivating adjustment for local gray-matter volume (GMV) and total intracranial volume (TIV) when characterizing functional associations.

We addressed three linked questions in 101 youths aged 8–18 years. First, which intrinsic signal properties show age-related variation across large-scale cortical networks? Second, does age shape their associations with contextual control, action cancellation, interference resolution, and oculomotor suppression? Third, do the spatial distributions of these associations follow established dimensions of cortical organization? We analyzed raw and within-brain spatially standardized BEN, ALFF, fALFF, and ReHo using voxelwise and seven-network models containing linear, quadratic, and cubic age terms. Local GMV, TIV, sex, and head motion were included to characterize functional associations conditional on observed anatomy and motion. Spatial correspondence with the S–A axis and three reference connectivity gradients connected local effects to broader cortical organization. Full-range analyses were primary, and age trimming evaluated sensitivity to the youngest and oldest participants.

## 2. Materials and methods

### 2.1 Participants and study design

Data came from the youth inhibitory-control cohort described by Cardinale et al. (2024), publicly available through OpenNeuro dataset ds005754, version 1.1.1 (Cardinale et al., 2026; https://doi.org/10.18112/openneuro.ds005754.v1.1.1). The release includes structural and resting-state MRI, task behavior, and phenotypic data. All participants in the local imaging pool were represented in this release with T1-weighted and three-echo resting-state files. Existing imaging derivatives and the original, unrounded age values were retained for the present analyses; release provenance is documented in Supplementary Section S1. This study was a secondary analysis of deidentified public data; the original study was approved by the NIH Institutional Review Board. The analysis was cross-sectional. Of 104 participants with complete matched functional metrics, three (sub-23628, sub-23807, and sub-24276) were excluded because severe tissue-segmentation failure mislabeled substantial gray matter as cerebrospinal fluid. This failure invalidated their GMV maps and could also contaminate tissue-derived nuisance signals. The primary sample therefore comprised 101 participants aged 8.10–17.98 years. Task-specific maximum samples were retained because task availability differed across participants (Table 1).

**Table 1.** Task-specific sample sizes.

| **Outcome** | **Primary n** | **Age-trimmed n** |
|---|---|---|
| AX-CPT | 53 | 50 |
| Anti-saccade | 48 | 41 |
| Stop-signal | 77 | 71 |
| Flanker | 91 | 83 |
| Composite | 59 | 54 |

### 2.2 MRI data acquisition

The source protocol acquired resting-state functional images while participants viewed a white fixation cross on a black background for 12 min. A three-echo echo-planar imaging sequence used a repetition time of 2,000 ms, nominal echo times of 14.8, 28.4, and 42.0 ms reported in the source protocol, a flip angle of 77°, a 64 × 64 matrix, and 34 interleaved axial slices of 3.8-mm thickness. T1-weighted structural images were acquired with a magnetization-prepared rapid gradient-echo sequence at 1-mm isotropic resolution and a flip angle of 7° (Cardinale et al., 2024; OpenNeuro ds005754). The local fMRIPrep metadata identify all three echo images as sources of the functional derivative and confirm a 2-s repetition time. For the present analyses, four initial volumes were removed and a common 300-volume segment (10 min) was retained for every functional metric.

### 2.3 Resting-state preprocessing

Resting-state data had been preprocessed with fMRIPrep 24.0.1 (Esteban et al., 2019) and analyzed in MNI152NLin6Asym 2-mm space. The first four volumes were excluded, and all functional metrics were computed from an identical 300-volume segment (TR=2.0 s; 10 min). Nuisance regression included white-matter and cerebrospinal-fluid signals, standardized DVARS, framewise displacement, RMS displacement, temporal CompCor components and three anatomical CompCor components (Behzadi et al., 2007), and six rigid-body motion parameters. Global signal regression and volume censoring were not used. Mean framewise displacement was retained as a model covariate because head motion can bias developmental resting-state estimates (Satterthwaite et al., 2012, 2013).

### 2.4 BOLD metrics

BEN was estimated using sample entropy (Richman and Moorman, 2000) as implemented in BENtbx (Wang et al., 2014). Time series were detrended, nuisance-regressed, filtered at 0.009–0.08 Hz, and standardized. Sample entropy used embedding dimension m=3 and tolerance r=0.60 times the time-series standard deviation. ALFF was calculated as the mean Fourier amplitude within 0.009–0.08 Hz (Zang et al., 2007). fALFF was calculated as the sum of amplitudes within that band divided by the sum across all positive frequencies (Zou et al., 2008). Both amplitude measures were estimated after 6-mm FWHM presmoothing. ReHo was calculated using Kendall’s coefficient of concordance within 27-voxel neighborhoods from band-pass-filtered data before spatial smoothing (Zang et al., 2004). Final metric maps were smoothed with an 8-mm FWHM kernel. For each metric, raw and within-brain spatial-z maps were analyzed; spatial-z maps describe relative regional expression.

### 2.5 Gray-matter volume

Native-space gray-matter probability maps from fMRIPrep were processed with an FSL-VBM workflow (Douaud et al., 2007; Smith et al., 2004) using a study-specific template, nonlinear FNIRT registration, Jacobian modulation, and 8-mm FWHM smoothing. Maps were regridded to the functional analysis space. Primary voxelwise models included the local voxelwise GMV value and TIV; network models included the corresponding network-average GMV and TIV. The three segmentation-failure participants were excluded from all functional analyses rather than selectively omitted from the structural covariates. The use of a hybrid fMRIPrep–FSL workflow is supported by comparative evidence of strong age-prediction performance alongside established alternatives such as CAT (Antonopoulos et al., 2023).

### 2.6 Behavioral measures

Task behavior was reconstructed from the public trial-level files accompanying Cardinale et al. (2024). AX-CPT contextual performance was quantified as the proportion of correct AX responses minus the proportion of incorrect BX responses. This accuracy-difference score was used in the present analyses; it is distinct from conventional signal-detection d′ because normal-quantile transforms were not applied. Anti-saccade performance was the proportion of all trials scored as correct. Stop-signal reaction time was estimated by the mean method as mean correct-go reaction time minus mean stop-signal delay; a 0.5-s pre-go interval was subtracted from the exported delay values before estimation. This estimator differs from the integration method recommended in contemporary guidance (Verbruggen et al., 2019). Flanker interference was the difference in mean reaction time between correct incongruent and congruent trials. Stop-signal and Flanker values were sign-reversed before standardization so that higher scores indicated better performance for each task. The composite was the mean of available standardized task scores and required at least three quality-controlled tasks. Task-specific analyses retained their maximum eligible samples; the composite was available for 59 participants.

### 2.7 Statistical analysis

Age was standardized and represented by linear, quadratic, and cubic terms. For each network, the primary age statistic was the joint 3-degree-of-freedom test of these terms; this tests an overall age association and does not by itself establish curvature beyond a linear trend. A separate 2-degree-of-freedom test of the quadratic and cubic terms evaluated departure from linearity. Task models included task performance, the three age terms, task interactions with each age term, sex, mean framewise displacement, TIV, and either local voxelwise or network GMV. The primary developmental statistic was a joint 3-degree-of-freedom test of task×age, task×age^2^, and task×age^3^, evaluating whether the task association varied across the modeled age function.

Voxelwise inference used 5,000 Freedman–Lane permutations (Winkler et al., 2014), a cluster-forming threshold of p<.001, and cluster-mass FDR at q<.05. FDR was applied separately within each metric-by-task map; there was no additional correction across tasks, metrics, or raw versus spatial-z scales. Seven-network analyses used Schaefer-200 parcels (Schaefer et al., 2018) assigned to the Yeo seven-network system (Yeo et al., 2011), heteroskedasticity-consistent HC3 standard errors (MacKinnon and White, 1985), and Benjamini–Hochberg FDR (Benjamini and Hochberg, 1995) across the seven networks within each metric-by-task-by-scale family. The same family definition was used for omnibus age associations.

Sensitivity analyses repeated the GMV-adjusted models after trimming the youngest and oldest 5% of the primary sample (age 8.94–17.45 years; n=91) and repeated the full-sample analysis without GMV and TIV. The full age range and GMV-adjusted model were designated as primary. Trimmed-only or GMV-sensitive findings were interpreted as sensitivity results. Full sensitivity results are reported in Tables S1, S3–S4, and S6 and Figure S1.

### 2.8 Cortical hierarchy analyses

We related unthresholded parcel-level joint age and task-by-age Wald statistics to the sensorimotor–association (S–A) axis (Sydnor et al., 2021) and the first three Margulies et al. (2016) connectivity gradients distributed through neuromaps (Markello et al., 2022). Functional and GMV values were averaged in Schaefer-400 parcels (Schaefer et al., 2018) with 17-network assignments (Yeo et al., 2011) within the fixed functional mask; 390 parcels had coverage. HC3 models included the same age terms and covariates as the network analyses, with matching parcel GMV. Spearman correlations were assessed with 5,000 paired hemispheric spherical rotations and one-to-one parcel reassignment (Alexander-Bloch et al., 2018). Positive correlations indicate stronger joint-effect evidence at higher reference values. The S–A age analyses report FDR across four raw metrics and separately across four spatial-z metrics; S–A task-wise results are reported as individual spin-test p values without pooling across tasks or metrics. Gradient analyses report BH-FDR across Gradients 1–3 within each metric-by-task family, with no pooling across tasks or metrics. Complete test families are reported in Supplementary Tables S9–S11.

## 3. Results

### 3.1 Age-related organization of intrinsic metrics

After local GMV and TIV adjustment, spatial-z ALFF showed corrected age associations under the cubic model in the somatomotor network (Wald χ^2^=13.56, q=.018) and salience/ventral-attention network (χ^2^=12.77, q=.018; Figure 1). No raw metric or other spatial-z metric survived seven-network FDR in the full sample. Separate tests of curvature beyond a linear age term did not survive seven-network FDR (Figure S3). The supported pattern therefore concerns age-related variation in the relative spatial distribution of ALFF, not a uniform increase or decrease in absolute ALFF. Remaining spatial-z network trajectories are shown in Figure S2, and significant omnibus age results are tabulated in Table S7.

**Figure 1.**
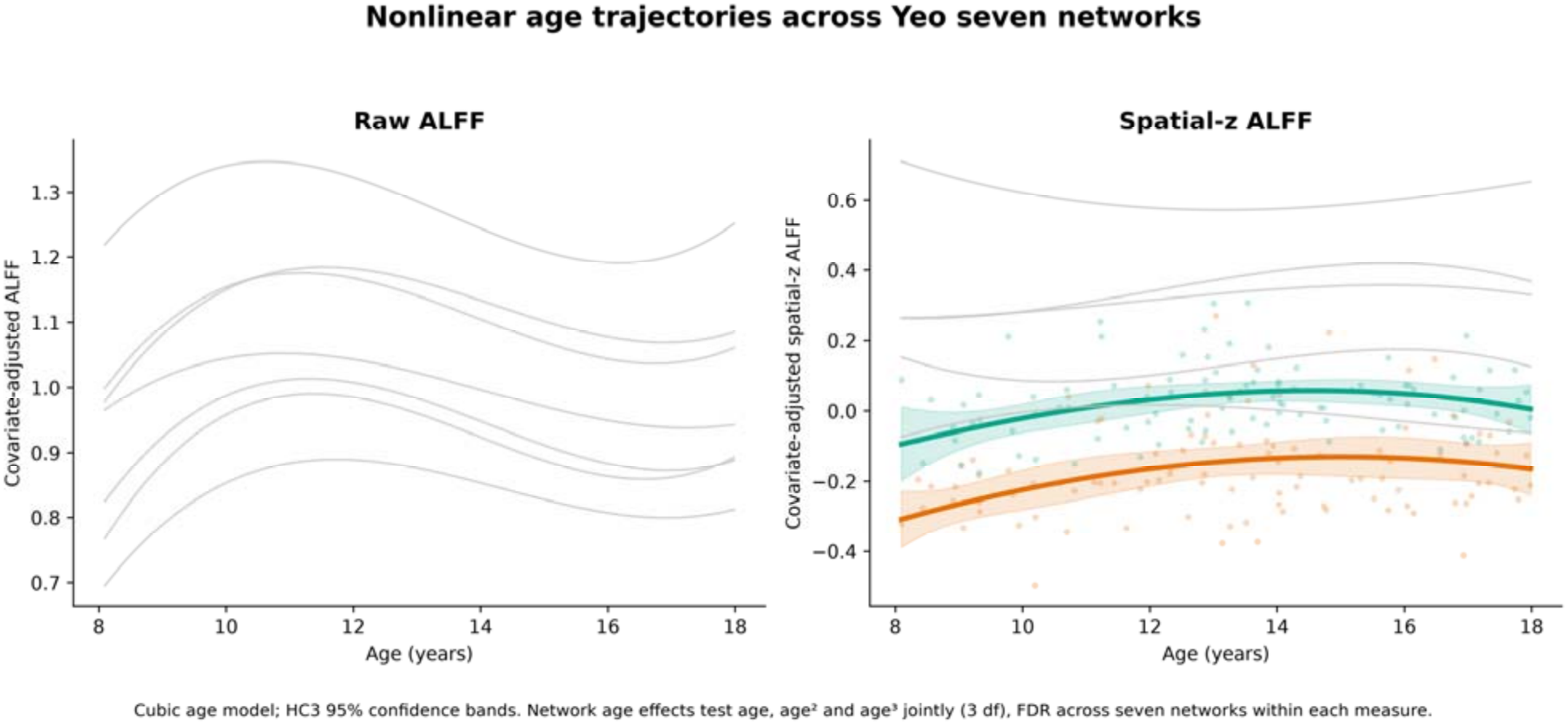
Raw ALFF (left) and within-brain spatial-z ALFF (right) age associations across seven networks. Colored curves and points identify networks with a significant joint cubic age association after FDR across seven networks; gray curves show the remaining networks. The omnibus HC3 Wald test evaluates age, age^2^, and age^3^ jointly (3 df); it does not alone test curvature beyond a linear trend. Only spatial-z ALFF was significant: somatomotor and salience/ventral-attention networks (both q=.018). Shading shows pointwise 95% confidence bands.

### 3.2 Voxelwise task-by-age interactions

Eight metric-by-task families contained at least one cluster surviving cluster-mass FDR (Table 2; Figure 2). AX-CPT performance showed the broadest pattern: corrected interactions were detected for spatial-z BEN, ALFF, fALFF, and ReHo. Spatial-z ALFF also interacted with Flanker performance. Anti-saccade performance interacted with raw fALFF and spatial-z ALFF, and composite performance interacted with raw fALFF. Raw BEN, raw ALFF, and raw ReHo produced no corrected voxelwise interactions. The cross-metric AX-CPT pattern indicates that contextual control was related to age-dependent spatial organization across several BOLD signal properties. Complete cluster coordinates are in Table S2; non-AX-CPT maps are shown in Figure S4.

**Table 2.** Primary GMV-adjusted voxelwise task-by-age interactions.

| **Metric** | **Task** | **n** | **Clusters** | **Cluster sizes** | **q range** |
|---|---|---|---|---|---|
| fALFF | Anti-saccade | 48 | 2 | 265, 338 | .032 |
| fALFF | Composite | 59 | 1 | 378 | .050 |
| zBEN | AX-CPT | 53 | 2 | 204, 224 | .030 |
| zALFF | AX-CPT | 53 | 3 | 281, 571, 220 | 0.016–0.035 |
| zALFF | Anti-saccade | 48 | 2 | 296, 410 | .021 |
| zALFF | Flanker | 91 | 1 | 324 | .048 |
| zfALFF | AX-CPT | 53 | 2 | 213, 375 | 0.020–0.041 |
| zReHo | AX-CPT | 53 | 1 | 217 | .016 |

**Figure 2.**
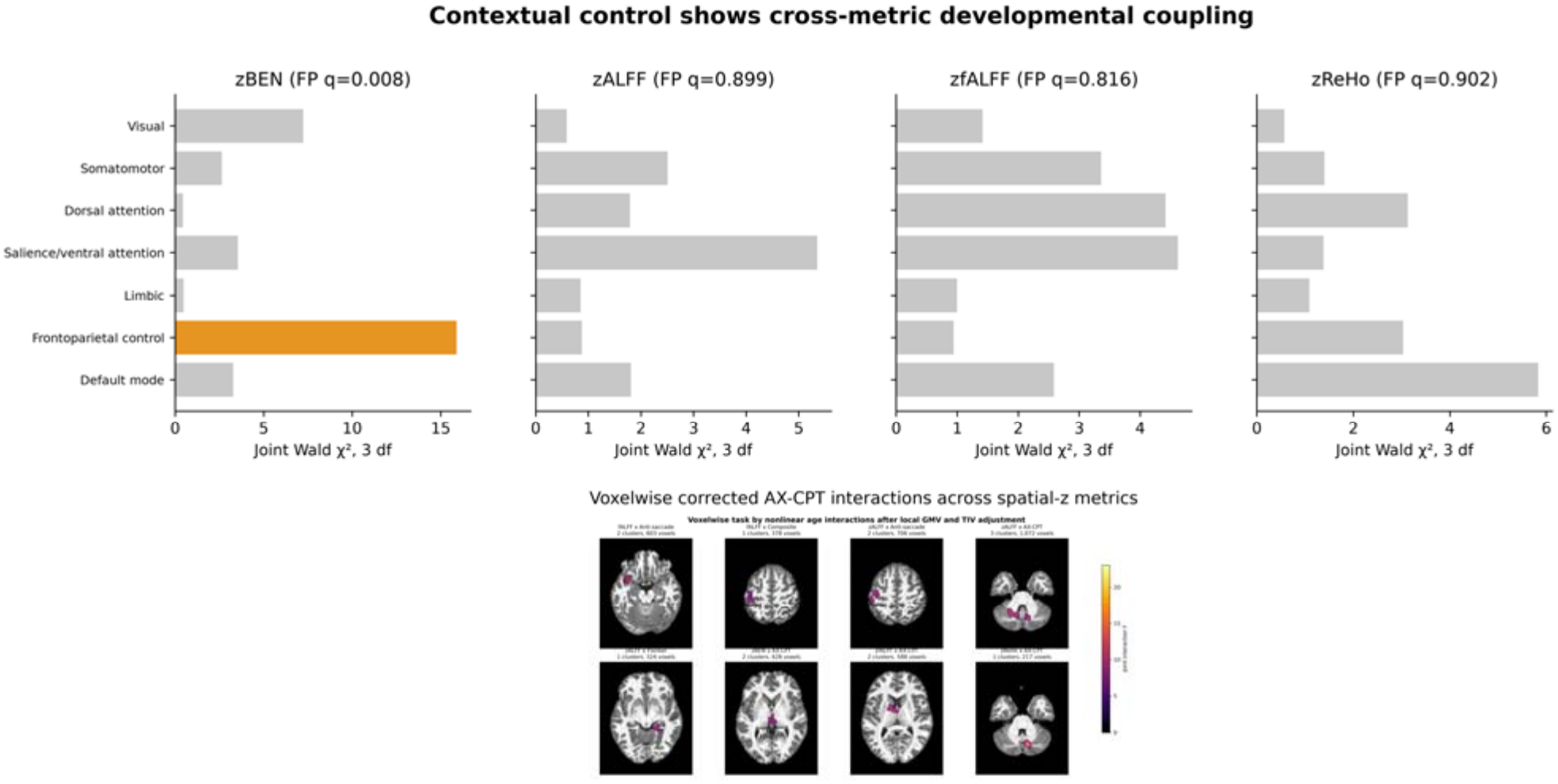
AX-CPT task-by-age interactions across spatial-z metrics. Top: seven-network joint Wald statistics, with color indicating network-level FDR q<.05. Bottom: voxelwise observed F maps restricted to clusters surviving permutation cluster-mass FDR within each metric-by-task map. The joint tests include task interactions with linear, quadratic, and cubic age terms; the F and Wald statistics are unsigned.

### 3.3 Network-level task associations

Network analyses separated task main effects from developmental interactions. Spatial-z BEN showed a frontoparietal AX-CPT interaction (χ^2^=15.90, q=.008), directly aligning the cross-metric voxelwise AX-CPT result with a canonical control network. Stop-signal performance was positively associated with frontoparietal BEN in both raw (β=.309, q=.028) and spatial-z form (β=.409, q<.001), without a corrected age interaction. Thus, contextual control was linked to age-dependent frontoparietal BEN organization, whereas action cancellation was linked to a more age-general frontoparietal BEN difference. Complete significant network results are in Table S5; the conditional frontoparietal BEN slope is illustrated in Figure S5.

Other corrected network findings were metric and task specific. Anti-saccade performance showed raw ReHo main effects in visual, dorsal-attention, salience/ventral-attention, and frontoparietal networks; raw fALFF showed a limbic developmental interaction; and spatial-z ReHo showed a limbic main effect. Spatial-z ALFF showed main effects for AX-CPT in the somatomotor network and anti-saccade in the limbic network. The composite score showed a raw fALFF interaction in the somatomotor network. The limited overlap among these findings is consistent with differentiated, rather than interchangeable, intrinsic signal properties.

### 3.4 Sensitivity analyses

Age trimming retained 91 participants overall and 41–83 participants across tasks. Five primary voxelwise families remained significant: spatial-z ALFF with AX-CPT and Flanker, and spatial-z BEN, fALFF, and ReHo with AX-CPT. This stable core supports the interpretation that AX-CPT developmental coupling was not driven solely by the age endpoints. Spatial-z BEN, fALFF, and ReHo Flanker effects emerged only after trimming and were treated as sensitivity findings. Raw fALFF anti-saccade/composite and spatial-z ALFF anti-saccade findings did not remain corrected after trimming.

The frontoparietal spatial-z BEN AX-CPT interaction and the spatial-z BEN stop-signal main effect remained significant after age trimming. The full-sample analysis without GMV and TIV reproduced several major patterns, including spatial-z ALFF, fALFF, and ReHo AX-CPT interactions, but the spatial-z BEN AX-CPT voxelwise interaction was not corrected in that model. This difference suggests that structural adjustment altered functional variance rather than revealing a simple structure-free effect; accordingly, the GMV-adjusted model remains primary and the no-GMV results are reported only as a supplement.

### 3.5 Structural results

GMV itself showed strong nonlinear age effects across all seven networks after TIV, sex, and motion adjustment. In contrast, neither voxelwise nor network-level GMV task main effects or task-by-age interactions survived correction. The functional task findings therefore were not mirrored by corrected structural task associations, although cross-sectional covariate adjustment cannot establish functional independence from maturational anatomy. See Table S8 and Figure S5 for network GMV results.

### 3.6 Cortical hierarchy of age and control-related effects

Raw BEN age-effect evidence was stronger toward association cortex on the S–A axis (ρ=.325, spin p=.0002; age-family q=.0008) and Gradient 1 (ρ=.364, q=.0006). The spatial-z ALFF × AX-CPT × age map showed the opposite spatial profile, with stronger joint-effect evidence toward sensorimotor cortex on both the S–A axis (ρ=−.227, spin p=.0106) and Gradient 1 (ρ=−.283, q=.0024). This convergence across independent references links age-related BEN organization and age-dependent contextual-control coupling to different positions along the cortical hierarchy (Figures 3 and 4). Figure 3 shows the reference surfaces and the unthresholded 400-parcel zALFF interaction map; Figure 4 summarizes the tested spatial correspondences.

**Figure 3.**
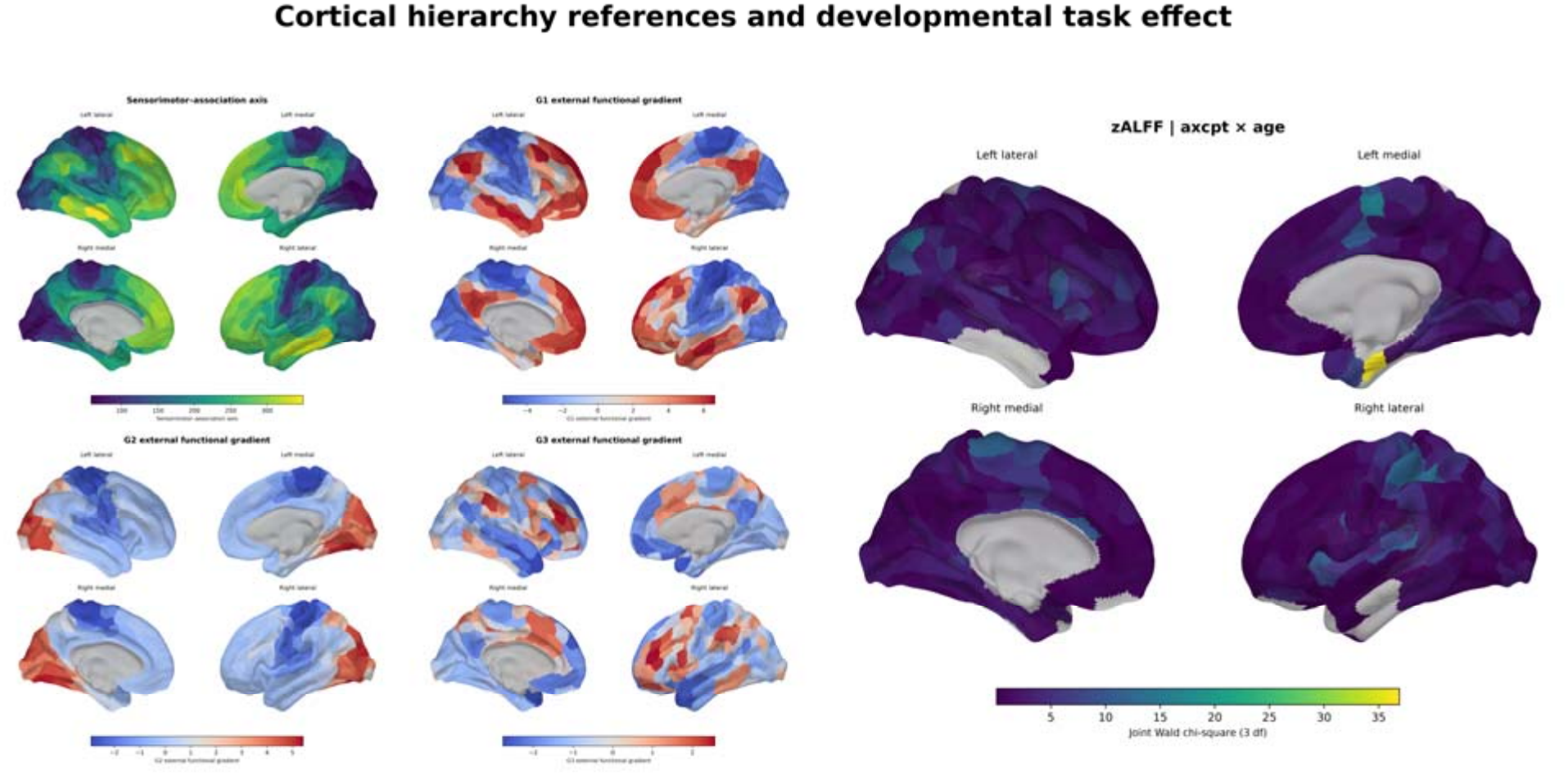
Cortical hierarchy references and the AX-CPT effect map. Left: external sensorimotor–association (S–A) axis and Gradients 1–3. Right: unthresholded 400-parcel spatial-z ALFF × AX-CPT × age joint Wald statistic. The parcelwise correlations with these references are summarized in Figure 4 across 390 parcels with functional-mask coverage. The map shows unsigned evidence, not parcelwise corrected significance.

**Figure 4.**
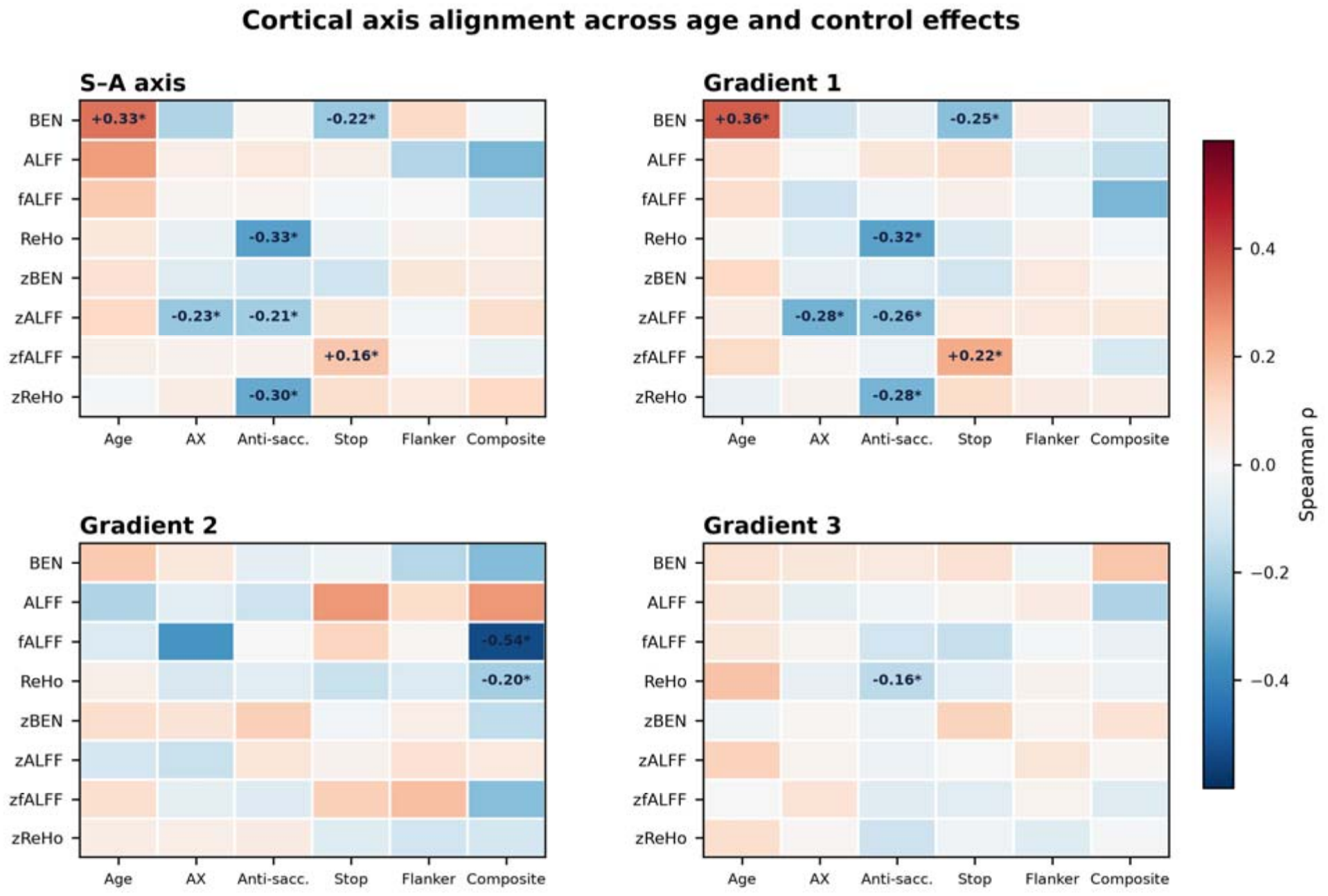
Cortical hierarchy of age and task-by-age effects. Colors show Spearman correlations between external S–A or Gradient 1–3 reference values and unthresholded joint Wald statistics across 390 covered parcels; significant cells are annotated with ρ. Task labels: AX, AX-CPT; Anti-sacc., anti-saccade; Stop, stop-signal. S–A task cells use individual two-sided spin p<.05 and age cells use within-scale q<.05. Gradient cells use q<.05 after FDR across three gradients within each metric-by-task family. Full results are in Tables S9–S11.

Additional task-specific associations extended beyond the S–A/Gradient 1 continuum: Gradient 2 aligned with raw fALFF and ReHo composite interactions, and Gradient 3 aligned with the raw ReHo anti-saccade interaction. Gradient 1 also aligned with stopping-related BEN and spatial-z fALFF, and with anti-saccade ReHo, spatial-z ALFF, and spatial-z ReHo. The S–A axis showed additional individual spin-test associations. These patterns support differentiated spatial organization across control demands; full estimates and all tested relationships are reported once in Supplementary Tables S9–S11. Correlations summarize alignment of unthresholded joint statistics across covered parcels and do not establish parcelwise significance or age-related change in the reference gradients.

## 4. Discussion

The principal finding is that the relationship between intrinsic brain activity and inhibitory control varies with both age and cognitive demand. Contextual performance on the AX-CPT showed the broadest voxelwise age interactions across spatially standardized BEN, ALFF, fALFF, and ReHo, together with a frontoparietal BEN interaction. Better action cancellation was associated with higher frontoparietal BEN across the sampled age range. These behavioral associations were embedded within a broader spatial organization: network-level ALFF age associations involved somatomotor and salience/ventral-attention systems, while age and task-interaction maps aligned with different cortical reference axes. Together, the findings suggest that developing control is associated with a changing relationship between specific cognitive demands and the intrinsic properties of distributed brain systems.

### 4.1 Contextual control links intrinsic activity to developing goal-directed behavior

Contextual control depends on carrying information forward and using it to guide a later response. The AX-CPT therefore places demand on goal maintenance, cue interpretation, and the selection of an appropriate action. The dual-mechanisms framework links sustained goal maintenance to anticipatory control that biases subsequent processing (Braver, 2012). Within this framework, the broad AX-CPT pattern suggests that age differences in contextual performance are accompanied by differences in how several properties of spontaneous activity relate to control. The frontoparietal BEN interaction gives this distributed pattern a connection to a canonical control system. Its persistence at the network level after age trimming further supports the relevance of frontoparietal temporal irregularity across the central age range.

This interpretation fits developmental accounts in which executive function improves through the coordinated maturation of specialized systems. Keller et al. (2023) describe attention and frontoparietal systems as integrative components linking hierarchically organized processing systems. The present results add local signal properties to that account. Contextual performance was associated with age-dependent variation in temporal irregularity, fluctuation amplitude, spectral composition, and local synchrony, suggesting several measurable dimensions through which intrinsic organization can relate to the demands of maintaining and applying context. Action cancellation showed a different profile: higher frontoparietal BEN accompanied better stopping without a corrected network age interaction. This contrast highlights the value of retaining individual control tasks when studying the developing brain.

The frontoparietal BEN findings also connect with evidence linking temporal irregularity to motivational traits. In young adult women, progesterone was negatively associated with BEN in dorsolateral prefrontal and posterior parietal regions, while left dorsolateral prefrontal BEN was positively associated with BAS-drive (Song and Wang, 2025). Together with the present task-based findings, this work identifies frontoparietal temporal irregularity as a candidate feature linking intrinsic activity to different aspects of goal-directed behavior. Its behavioral relevance appears to depend on the demands being measured, spanning motivational drive, contextual performance, and action cancellation.

### 4.2 Complementary signal properties reveal different aspects of developmental organization

The cross-metric AX-CPT pattern is informative because the four measures characterize distinct features of BOLD activity. BEN describes temporal irregularity, ALFF fluctuation amplitude, fALFF relatively low-frequency spectral content, and ReHo local temporal concordance. Previous work found that BEN–fALFF associations vary across regions, including positive and negative relationships (Song et al., 2019). Pharmacological evidence adds a complementary observation: oxytocin increased temporoparietal BEN in younger adults and decreased it in older adults, whereas fALFF increased in the same region, particularly in younger adults (Song and Wang, 2026). These results establish an empirical basis for examining temporal and spectral properties together. In the present cohort, their distinct spatial associations suggest that a shared control demand can relate to different features of intrinsic activity across different cortical locations.

BEN also differentiated age-related organization from behavioral coupling. Raw BEN age-effect evidence was stronger toward association cortex, whereas frontoparietal BEN was linked to contextual performance and action cancellation. An earlier movie-viewing preprint similarly reported spatially selective BEN–age associations in children despite no significant within-childhood age association in global mean BEN (Song, 2025). Viewed together, these studies motivate a regional account of developmental temporal irregularity: its distribution across systems, and its relationship to cognition, can carry information that is obscured by a whole-brain average. The current study extends this question from naturalistic viewing to resting-state associations with specific control demands.

### 4.3 Cortical hierarchy organizes distinct age and control associations

The spatial analyses place the local findings within a broader developmental framework. Functional-connectivity gradients distinguish sensory and motor systems from transmodal association cortex (Margulies et al., 2016), and their organization changes across childhood and adolescence (Dong et al., 2021). In four independent developmental datasets, Luo et al. (2024) showed that connectivity development follows the sensorimotor–association axis and enhances the differentiation of cortical connectivity profiles. The present findings extend the relevance of this spatial framework to local BOLD signal properties. Raw BEN age-effect evidence was stronger toward the association end of both the S–A axis and Gradient 1, locating age-related variation in temporal irregularity within systems characterized by prolonged functional development (Sydnor et al., 2021).

Contextual-control coupling showed a complementary spatial organization. The spatial-z ALFF × AX-CPT × age map aligned toward the sensorimotor end of the S–A axis and Gradient 1. Alongside the frontoparietal BEN interaction, this finding links contextual performance to both a control-network property and distributed variation in relative amplitude across lower-order systems. A plausible interpretation is that development changes how control-relevant intrinsic organization is expressed across systems involved in maintaining goals and implementing responses. This interpretation connects the findings to theories of multisystem coordination while retaining a distinction between the regional expression of signal properties and the connectivity architecture studied in prior work.

Other control measures extended the spatial pattern beyond the principal sensorimotor–association continuum. Gradient 2 aligned with raw fALFF and ReHo composite-interaction maps, with stronger evidence toward its somatomotor end. Gradient 3 aligned with the raw ReHo anti-saccade interaction map. These findings indicate that different control demands relate to complementary dimensions of cortical organization. The S–A axis and Gradient 1 provide a reference for broad hierarchical differentiation, while additional gradients capture spatial distinctions relevant to sensory and action-related systems. This multidimensional organization offers a more specific account of intrinsic control associations than a single common anatomical signature.

### 4.4 Relative functional expression develops within a changing anatomical landscape

The predominance of spatially standardized metrics in the AX-CPT findings emphasizes the relative organization of intrinsic activity within each brain. Spatial-z measures compare regional values with an individual’s whole-brain distribution, making them sensitive to differences in where a signal property is expressed most strongly. The network age associations in spatial-z ALFF, together with the spatial-z task interactions, suggest that relative regional expression is a useful level for characterizing developmental brain–behavior relationships. Raw BEN contributes a complementary view through its age-effect alignment with association cortex. These observations link development to several levels of intrinsic organization, spanning original metric values, relative topography, and behavioral associations.

This functional organization unfolds alongside substantial anatomical maturation. GMV showed age associations across all seven networks, consistent with the broad nonlinear anatomical trajectories described by lifespan brain charts (Bethlehem et al., 2022). Local GMV and TIV adjustment allowed the primary analyses to characterize functional associations within this changing structural landscape. The presence of core AX-CPT findings after adjustment and age trimming strengthens their relevance to intrinsic functional organization, while the sensitivity of voxelwise BEN results to anatomical adjustment highlights the close relationship between the structural and functional features of developing cortex.

### 4.5 Scope and future directions

The findings identify age-related differences in brain–behavior relationships in a cross-sectional cohort. Longitudinal data are needed to determine how these relationships change within individuals and whether they accompany gains in contextual control or action cancellation. Task-specific samples varied in size and availability, and the reconstructed AX-CPT accuracy-difference score captures contextual performance rather than a process-pure measure of proactive control. Spatial correlations with joint age statistics localize the strength of age-related evidence; directional trajectories require examination of the underlying fitted relationships. Adult reference gradients provide a common cortical coordinate system, and individual developmental gradients could further clarify how local signal properties and connectivity organization evolve together. Inference is defined within each prespecified metric-by-task correction family. These considerations motivate replication with larger, longitudinal samples and harmonized behavioral measures.

The resulting developmental account centers on the changing correspondence between intrinsic activity and specific control demands. Contextual control showed broad age-dependent coupling across complementary signal properties, action cancellation was associated with frontoparietal temporal irregularity, and their spatial distributions occupied different cortical organizing dimensions. This framework connects local BOLD dynamics to the differentiation of cognitive control during adolescence and provides testable targets for studies of individual developmental trajectories.

## 5. Conclusion

Distinct control demands showed different age-dependent relationships with intrinsic brain activity. Contextual control had the broadest cross-metric voxelwise pattern and a frontoparietal BEN interaction, whereas action cancellation was associated with frontoparietal BEN across the sampled ages. Across cortical hierarchy, age and task-related effects occupied different spatial positions. These findings place complementary intrinsic signal properties within a differentiated account of control development, connecting component-specific behavior to the organization of cortical systems.

## Data and code availability

The source cohort is publicly available through OpenNeuro dataset ds005754, version 1.1.1 (https://doi.org/10.18112/openneuro.ds005754.v1.1.1).

