## Supplementary Materials for "Distinct developmental organization of intrinsic brain dynamics supports different components of inhibitory control during adolescence"

### Supplementary overview

These materials document sample derivation, preprocessing, statistical families, complete corrected cluster coordinates, seven-network findings, age-range sensitivity analyses, and matched-sample models without GMV/TIV. The primary estimand is the N=101 full-age-range functional association conditional on local GMV and TIV.

### S1. Sample derivation and quality control

Dataset provenance. The current public release is OpenNeuro ds005754 v1.1.1. Its repository contains T1-weighted and three-echo resting-state MRI files for all 104 participants in the local imaging pool. The existing fMRIPrep derivatives identify ds004935 v1.0.1 as their historical source; behavioral and phenotypic files were retrieved from ds005754 v1.1.1 and matched by participant identifier. The original unrounded ages were retained, because the release notes report rounding age to one decimal place in version 1.1.0. The old public repository was unavailable at verification, so identity of the old and current raw imaging files could not be checked by checksum; existing derivatives were retained rather than described as newly processed from v1.1.1. Acquisition parameters in the main text follow the published source protocol. Some example BIDS sidecars differ from the protocol summary (including structural inversion time and echo-time labels); unverified sequence-specific values were therefore omitted rather than generalized across the cohort.

The functional metric set initially contained 104 participants. Three participants were excluded from all analyses because visual and quantitative inspection showed severe tissue-classification failure: sub-23628, sub-23807, and sub-24276. In each case, gray matter was substantially mislabeled as cerebrospinal fluid. Because tissue masks contributed both to voxelwise GMV and to tissue-derived nuisance regressors, selective structural exclusion would not have resolved the quality problem. The final primary sample was N=101.

**Table S1. Sample sizes by analysis.**

| **Analysis** | **Overall n** | **AX-CPT** | **Anti-saccade** | **Stop-signal** | **Flanker** | **Composite** |
| --- | --- | --- | --- | --- | --- | --- |
| Primary GMV+TIV | 101 | 53 | 48 | 77 | 91 | 59 |
| Age 5th–95th percentile | 91 | 50 | 41 | 71 | 83 | 54 |
| Matched no GMV/TIV | 101 | 53 | 48 | 77 | 91 | 59 |

### S2. Image processing details

#### S2.1 Functional metrics

All functional measures used the same 300-volume segment after removal of four initial volumes. BEN used detrended, nuisance-regressed, 0.009–0.08-Hz filtered, standardized time series with Sample Entropy m=3 and r=0.60×SD. ALFF and fALFF used 6-mm presmoothing before metric estimation. ReHo was estimated before smoothing from band-pass-filtered data. Final maps for all four metrics were smoothed to 8-mm FWHM. Raw maps and within-brain spatial-z maps were retained as different estimands.

#### S2.2 Structural processing

Gray-matter probability maps were processed with a study-specific FSL-VBM workflow, nonlinear registration, Jacobian modulation, and 8-mm smoothing. TIV was included as a global structural covariate. Voxel models used local modulated GMV; network models used the matching network-average GMV.

Behavioral reconstruction. The analysis variable named axcpt_dprime_context in the scripts is AX accuracy minus BX error rate, without normal-quantile transformation. It is an accuracy-difference contextual-performance score, not conventional d′. Stop-signal reaction time uses the mean method with a 0.5-s offset removed from exported stop-signal delays. These definitions describe the existing analyzed variables; correcting the estimator would require new behavioral scores and reanalysis.

### S3. Statistical families and interpretation

The primary interaction was the joint 3-df test of task×age, task×age², and task×age³. Voxelwise tests used 5,000 Freedman–Lane permutations, cluster-forming p<.001, and cluster-mass FDR q<.05. Each metric×task map was one correction family. Network models used HC3 covariance and BH-FDR across seven networks within each metric×task×scale family. No correction was applied across tasks, metrics, or raw/spatial-z scales. Corrected inference applies within each declared family.

### S4. Primary voxelwise results

All clusters below survived permutation cluster-mass FDR in the full N=101 GMV+TIV-adjusted analysis. Peak statistics are joint F values for the three task-by-age interaction terms (linear, quadratic, and cubic age).

**Table S2. Significant primary voxelwise clusters.**

| **Metric** | **Task** | **n** | **ID** | **Voxels** | **Peak F** | **Peak MNI** | **q** |
| --- | --- | --- | --- | --- | --- | --- | --- |
| fALFF | Anti-saccade | 48 | 3 | 265 | 14.83 | (-36, 6, -24) | .032 |
| fALFF | Anti-saccade | 48 | 25 | 338 | 14.17 | (64, -26, 0) | .032 |
| fALFF | Composite | 59 | 3 | 378 | 9.00 | (-38, -32, 58) | .050 |
| zBEN | AX-CPT | 53 | 2 | 204 | 10.02 | (-36, -60, -32) | .030 |
| zBEN | AX-CPT | 53 | 13 | 224 | 11.96 | (-2, -22, 4) | .030 |
| zALFF | AX-CPT | 53 | 3 | 281 | 23.05 | (-34, -18, -22) | .018 |
| zALFF | AX-CPT | 53 | 5 | 571 | 11.59 | (-22, -54, -38) | .016 |
| zALFF | AX-CPT | 53 | 6 | 220 | 11.37 | (10, -66, -30) | .035 |
| zALFF | Anti-saccade | 48 | 1 | 296 | 11.02 | (-46, -34, 60) | .021 |
| zALFF | Anti-saccade | 48 | 10 | 410 | 10.18 | (34, -74, -30) | .021 |
| zALFF | Flanker | 91 | 17 | 324 | 9.76 | (20, -38, -8) | .048 |
| zfALFF | AX-CPT | 53 | 2 | 213 | 12.15 | (-14, -66, -30) | .041 |
| zfALFF | AX-CPT | 53 | 6 | 375 | 12.39 | (0, -4, 10) | .020 |
| zReHo | AX-CPT | 53 | 7 | 217 | 18.66 | (10, -70, -32) | .016 |

### S5. Age-range sensitivity

The 5th–95th percentile analysis retained ages 8.94–17.45 years (N=91). Results emerging only after trimming are sensitivity findings and are not promoted to primary evidence.

**Table S3. Significant GMV+TIV-adjusted voxelwise clusters after age trimming.**

| **Metric** | **Task** | **n** | **ID** | **Voxels** | **Peak F** | **Peak MNI** | **q** |
| --- | --- | --- | --- | --- | --- | --- | --- |
| zBEN | AX-CPT | 50 | 2 | 242 | 14.12 | (-34, -60, -32) | .031 |
| zBEN | AX-CPT | 50 | 16 | 203 | 11.65 | (-4, -22, 4) | .031 |
| zBEN | Flanker | 83 | 5 | 165 | 11.17 | (-14, 48, 24) | .043 |
| zBEN | Flanker | 83 | 12 | 227 | 10.77 | (20, -32, 34) | .043 |
| zALFF | AX-CPT | 50 | 4 | 264 | 22.31 | (-34, -18, -22) | .022 |
| zALFF | AX-CPT | 50 | 6 | 588 | 11.42 | (-24, -52, -38) | .013 |
| zALFF | AX-CPT | 50 | 8 | 212 | 11.40 | (10, -64, -30) | .038 |
| zALFF | Flanker | 83 | 17 | 669 | 10.69 | (34, -64, -28) | .008 |
| zfALFF | AX-CPT | 50 | 2 | 225 | 14.94 | (-14, -66, -30) | .029 |
| zfALFF | AX-CPT | 50 | 7 | 328 | 14.05 | (-12, 4, 8) | .025 |
| zfALFF | Flanker | 83 | 6 | 437 | 10.87 | (-26, -6, 62) | .007 |
| zfALFF | Flanker | 83 | 16 | 186 | 8.22 | (4, -50, -16) | .033 |
| zfALFF | Flanker | 83 | 18 | 191 | 11.97 | (8, -30, 4) | .033 |
| zfALFF | Flanker | 83 | 24 | 567 | 11.96 | (36, -22, 4) | .007 |
| zReHo | AX-CPT | 50 | 7 | 253 | 19.57 | (10, -70, -32) | .009 |
| zReHo | Flanker | 83 | 5 | 263 | 10.29 | (-20, 8, 28) | .017 |

### S6. Matched-sample models without GMV/TIV

These analyses used the same 101 participants and task-specific sample sizes as the primary analysis but omitted local GMV and TIV. They estimate functional associations without conditioning on observed anatomy and are provided as supplementary comparisons.

**Table S4. Significant voxelwise clusters without GMV/TIV.**

| **Metric** | **Task** | **n** | **ID** | **Voxels** | **Peak F** | **Peak MNI** | **q** |
| --- | --- | --- | --- | --- | --- | --- | --- |
| fALFF | Anti-saccade | 48 | 3 | 235 | 11.20 | (-34, 12, -18) | .044 |
| fALFF | Anti-saccade | 48 | 20 | 413 | 15.64 | (64, -26, 0) | .036 |
| fALFF | Composite | 59 | 1 | 411 | 9.52 | (-38, -32, 58) | .045 |
| fALFF | Composite | 59 | 3 | 211 | 9.87 | (-50, -6, 34) | .045 |
| fALFF | Composite | 59 | 22 | 224 | 10.70 | (56, -16, 10) | .045 |
| zBEN | Flanker | 91 | 2 | 212 | 9.02 | (-16, 50, 24) | .046 |
| zBEN | Flanker | 91 | 3 | 179 | 10.74 | (-14, -4, 40) | .046 |
| zALFF | AX-CPT | 53 | 4 | 261 | 18.28 | (-32, -18, -22) | .032 |
| zALFF | AX-CPT | 53 | 6 | 749 | 11.60 | (-20, -60, -32) | .014 |
| zALFF | AX-CPT | 53 | 7 | 267 | 10.90 | (10, -64, -30) | .032 |
| zALFF | Anti-saccade | 48 | 1 | 300 | 11.39 | (-52, -24, 54) | .030 |
| zALFF | Anti-saccade | 48 | 11 | 303 | 9.32 | (34, -76, -28) | .030 |
| zfALFF | AX-CPT | 53 | 1 | 259 | 12.46 | (-14, -68, -30) | .029 |
| zfALFF | AX-CPT | 53 | 6 | 253 | 10.55 | (-12, 0, 8) | .029 |
| zReHo | AX-CPT | 53 | 4 | 297 | 21.58 | (10, -70, -32) | .009 |

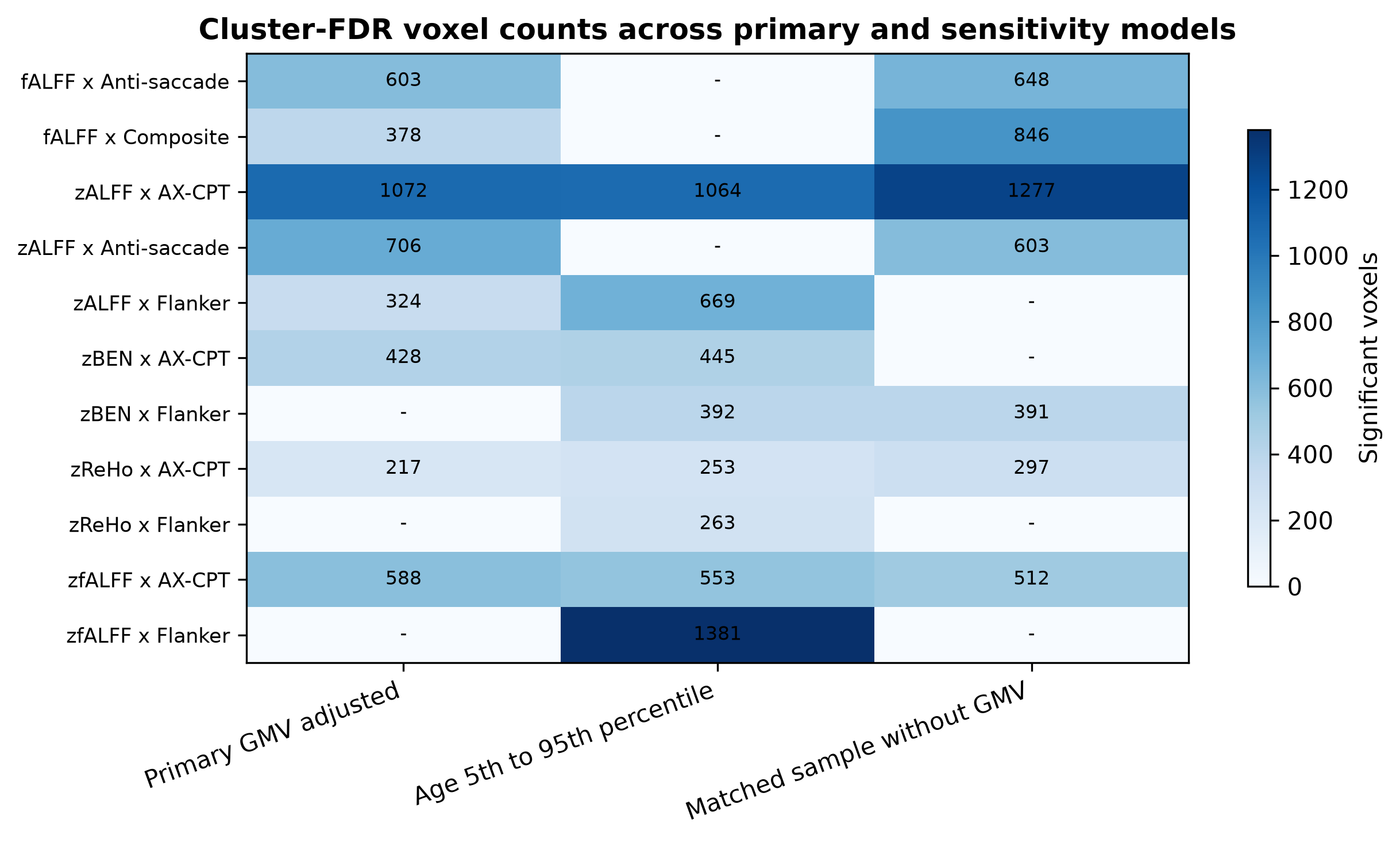

Figure S1. Numbers of voxels in corrected interaction clusters across the primary GMV-adjusted analysis, age-trimmed GMV-adjusted sensitivity analysis, and matched N=101 analysis without GMV/TIV. A dash indicates no surviving cluster; each cell summarizes one metric-by-task correction family.

### S7. Seven-network results

**Table S5. Significant primary functional network results (local GMV+TIV).**

| **Scale** | **Metric** | **Task** | **Network** | **n** | **Effect** | **Estimate** | **q** |
| --- | --- | --- | --- | --- | --- | --- | --- |
| raw | BEN | Stop-signal | Frontoparietal control | 77 | Task main | β=0.309 | .028 |
| raw | fALFF | Anti-saccade | Limbic | 48 | Task×age joint | χ²=15.46 | .010 |
| raw | fALFF | Composite | Somatomotor | 59 | Task×age joint | χ²=12.92 | .034 |
| raw | ReHo | Anti-saccade | Visual | 48 | Task main | β=0.406 | .026 |
| raw | ReHo | Anti-saccade | Dorsal attention | 48 | Task main | β=0.450 | .002 |
| raw | ReHo | Anti-saccade | Salience/ventral attention | 48 | Task main | β=0.313 | .036 |
| raw | ReHo | Anti-saccade | Frontoparietal control | 48 | Task main | β=0.259 | .036 |
| spatial_z | zBEN | AX-CPT | Frontoparietal control | 53 | Task×age joint | χ²=15.90 | .008 |
| spatial_z | zBEN | Stop-signal | Frontoparietal control | 77 | Task main | β=0.409 | <.001 |
| spatial_z | zALFF | AX-CPT | Somatomotor | 53 | Task main | β=-0.380 | .016 |
| spatial_z | zALFF | Anti-saccade | Limbic | 48 | Task main | β=-0.408 | .031 |
| spatial_z | zReHo | Anti-saccade | Limbic | 48 | Task main | β=-0.475 | .004 |

**Table S6. Significant age-trimmed functional network results (local GMV+TIV).**

| **Scale** | **Metric** | **Task** | **Network** | **n** | **Effect** | **Estimate** | **q** |
| --- | --- | --- | --- | --- | --- | --- | --- |
| raw | fALFF | Anti-saccade | Limbic | 41 | Task main | β=-0.343 | .047 |
| raw | fALFF | Anti-saccade | Frontoparietal control | 41 | Task×age joint | χ²=14.12 | .019 |
| raw | ReHo | Anti-saccade | Visual | 41 | Task main | β=0.408 | .019 |
| raw | ReHo | Anti-saccade | Dorsal attention | 41 | Task main | β=0.500 | <.001 |
| raw | ReHo | Anti-saccade | Salience/ventral attention | 41 | Task main | β=0.371 | .019 |
| raw | ReHo | Anti-saccade | Frontoparietal control | 41 | Task main | β=0.295 | .019 |
| spatial_z | zBEN | AX-CPT | Frontoparietal control | 50 | Task×age joint | χ²=15.25 | .011 |
| spatial_z | zBEN | AX-CPT | Default mode | 50 | Task main | β=-0.536 | .002 |
| spatial_z | zBEN | Stop-signal | Frontoparietal control | 71 | Task main | β=0.423 | <.001 |
| spatial_z | zBEN | Stop-signal | Default mode | 71 | Task main | β=0.329 | .012 |
| spatial_z | zALFF | AX-CPT | Somatomotor | 50 | Task main | β=-0.397 | .027 |
| spatial_z | zfALFF | Anti-saccade | Limbic | 41 | Task×age joint | χ²=16.96 | .005 |
| spatial_z | zReHo | Anti-saccade | Visual | 41 | Task×age joint | χ²=14.94 | .013 |
| spatial_z | zReHo | Anti-saccade | Limbic | 41 | Task main | β=-0.458 | .048 |

### S8. Nonlinear age effects

Table S7. Functional network omnibus age associations surviving FDR.

| **Sample** | **Scale** | **Metric** | **Network** | **n** | **χ²(3)** | **q** |
| --- | --- | --- | --- | --- | --- | --- |
| Primary | spatial_z | zALFF | Somatomotor | 101 | 13.56 | .018 |
| Primary | spatial_z | zALFF | Salience/ventral attention | 101 | 12.77 | .018 |
| Trimmed | spatial_z | zBEN | Default mode | 91 | 17.37 | .004 |
| Trimmed | spatial_z | zALFF | Salience/ventral attention | 91 | 12.19 | .024 |
| Trimmed | spatial_z | zALFF | Frontoparietal control | 91 | 16.91 | .005 |

Table S8. Network GMV omnibus age associations adjusted for TIV, sex, and motion.

| **Network** | **n** | **χ²(3)** | **q** |
| --- | --- | --- | --- |
| Visual | 101 | 35.45 | <.001 |
| Somatomotor | 101 | 23.37 | <.001 |
| Dorsal attention | 101 | 35.86 | <.001 |
| Salience/ventral attention | 101 | 43.13 | <.001 |
| Limbic | 101 | 21.69 | <.001 |
| Frontoparietal control | 101 | 40.46 | <.001 |
| Default mode | 101 | 46.37 | <.001 |

### S9. Null structural task results

No voxelwise GMV task-by-age interaction survived the cluster-based correction, and no seven-network GMV task main effect or joint nonlinear interaction survived seven-network FDR. These null corrected results are distinct from the robust age-related GMV trajectories in Table S8.

### S10. Reproducibility notes

- All primary and sensitivity voxelwise combinations completed without recorded failures.
- Observed maps and permutation outputs were retained for each metric-by-task family.
- The search mask and correction definition were held fixed within each comparison.
- BEN values were calculated from matched 300-volume time series with identical parameters.
- Raw and spatial-z results are reported separately because they answer different questions.

### S11. Cortical hierarchy analyses

Independent reference maps comprised the archetypal S–A axis of Sydnor et al. (2021) and adult group-average functional-connectivity Gradients 1–3 from Margulies et al. (2016), distributed by neuromaps (Markello et al., 2022). The S–A ranks were obtained in Schaefer-400 17-network order; gradients were averaged from matching fsLR 32k vertices into these parcels. Labels were matched by name. The volumetric Schaefer atlas was resampled to the functional grid using nearest-neighbor interpolation. Ten parcels had no functional-mask coverage and were excluded from correlations, leaving a fixed 390-parcel subset.

Raw and spatial-z subject maps were averaged within each covered parcel. BEN values were converted from the stored scale by dividing by 1000; spatial-z BEN was calculated within the same fixed whole-brain mask as the primary analyses before parcellation. Matching modulated GMV was averaged over identical parcel voxels. Age-only models included linear, quadratic and cubic standardized age, sex, mean framewise displacement, standardized TIV, and standardized parcel GMV. Task models additionally included standardized task performance and its three age interactions. The full cohort comprised 101 participants; task samples were AX-CPT n=53, anti-saccade n=48, stop-signal n=77, Flanker n=91, and composite n=59.

For each parcel, HC3 covariance estimates were used to obtain a joint 3-df Wald statistic for age terms or task-by-age interactions. Spearman correlations with reference values used all covered parcels, without voxel or parcel significance thresholding. These are parcel-level extensions of the primary analysis. Wald statistics are unsigned: positive spatial correlation indicates stronger statistical evidence toward the higher-value end of a reference map, not a positive age slope, better performance, or an increase in the underlying metric.

Spatial nulls used 5,000 random spherical rotations (seed 20261117). Spherical parcel centroids were calculated from the matching fsLR labels and spheres. Right-hemisphere rotations were mirrored relative to the left hemisphere; Hungarian assignment produced a one-to-one mapping within each hemisphere. The same rotations were used for all references and effects. Reference maps were rotated across 400 parcels and correlations recomputed in the fixed 390-parcel subset. Two-sided Monte Carlo p values were (1 + number of absolute null correlations at least as large as the observed correlation)/5001.

The S–A axis and connectivity gradients provide the spatial reference frameworks for interpreting age and task-by-age distributions. The S–A age analyses report BH-FDR across four raw metrics and separately across four spatial-z metrics. For the three connectivity gradients, BH-FDR was applied across Gradient 1–3 within each metric-by-task family (including age-only families). There was no pooling across tasks or metrics. All 48 S–A tests and 144 gradient tests, including null results, are reported below.

The released reference signs were retained. Gradient 1 had higher mean values in default-mode and control networks than visual and somatomotor networks; Gradient 2 had higher visual than somatomotor values; Gradient 3 had higher control/attention values than visual, somatomotor, and default-mode values. Thus, S–A and Gradient 1 describe related sensorimotor–association organization, while Gradients 2 and 3 provide additional spatial axes. These adult references were not estimated separately in the youth sample.

Validation reproduced 12 selected joint Wald statistics using an independent statsmodels HC3 implementation. All 5,000 spin assignments were verified as one-to-one hemisphere-preserving permutations. The observed zALFF–AX-CPT Spearman correlation was independently reproduced with scipy. Reference download URLs and SHA-256 checksums, parcel coverage, subject values, all effect maps in tabular form, and spin indices accompany the analysis code.

**Table S9. Spatial correspondence of joint age statistics with the S–A axis.**

| **Metric** | **n** | **ρ** | **Spin p** | **Age-family q** |
| --- | --- | --- | --- | --- |
| BEN | 101 | 0.325 | 0.0002 | 0.0008 |
| ALFF | 101 | 0.253 | 0.1618 | 0.2157 |
| fALFF | 101 | 0.159 | 0.1478 | 0.2157 |
| ReHo | 101 | 0.064 | 0.5449 | 0.5449 |
| zBEN | 101 | 0.087 | 0.2228 | 0.7782 |
| zALFF | 101 | 0.118 | 0.3891 | 0.7782 |
| zfALFF | 101 | 0.034 | 0.6959 | 0.9254 |
| zReHo | 101 | -0.008 | 0.9254 | 0.9254 |

**Table S10. All task-by-age spatial correspondence tests with the S–A axis.**

| **Metric** | **Task** | **n** | **ρ** | **Spin p** |
| --- | --- | --- | --- | --- |
| BEN | axcpt | 53 | -0.179 | 0.0870 |
| BEN | antisaccade | 48 | 0.010 | 0.9178 |
| BEN | stopsignal | 77 | -0.224 | 0.0010 |
| BEN | flanker | 91 | 0.115 | 0.2547 |
| BEN | composite | 59 | -0.008 | 0.9452 |
| ALFF | axcpt | 53 | 0.042 | 0.7187 |
| ALFF | antisaccade | 48 | 0.057 | 0.6763 |
| ALFF | stopsignal | 77 | 0.035 | 0.8124 |
| ALFF | flanker | 91 | -0.174 | 0.3729 |
| ALFF | composite | 59 | -0.274 | 0.2897 |
| fALFF | axcpt | 53 | 0.016 | 0.9286 |
| fALFF | antisaccade | 48 | 0.022 | 0.9100 |
| fALFF | stopsignal | 77 | -0.005 | 0.9606 |
| fALFF | flanker | 91 | 0.003 | 0.9808 |
| fALFF | composite | 59 | -0.118 | 0.5403 |
| ReHo | axcpt | 53 | -0.043 | 0.6109 |
| ReHo | antisaccade | 48 | -0.326 | 0.0006 |
| ReHo | stopsignal | 77 | -0.040 | 0.6521 |
| ReHo | flanker | 91 | 0.024 | 0.7698 |
| ReHo | composite | 59 | 0.041 | 0.6181 |
| zBEN | axcpt | 53 | -0.072 | 0.4671 |
| zBEN | antisaccade | 48 | -0.098 | 0.2294 |
| zBEN | stopsignal | 77 | -0.121 | 0.2164 |
| zBEN | flanker | 91 | 0.066 | 0.4101 |
| zBEN | composite | 59 | 0.055 | 0.5149 |
| zALFF | axcpt | 53 | -0.227 | 0.0106 |
| zALFF | antisaccade | 48 | -0.208 | 0.0218 |
| zALFF | stopsignal | 77 | 0.067 | 0.7143 |
| zALFF | flanker | 91 | -0.016 | 0.8836 |
| zALFF | composite | 59 | 0.095 | 0.3637 |
| zfALFF | axcpt | 53 | 0.025 | 0.8490 |
| zfALFF | antisaccade | 48 | 0.025 | 0.8302 |
| zfALFF | stopsignal | 77 | 0.162 | 0.0394 |
| zfALFF | flanker | 91 | -0.001 | 0.9910 |
| zfALFF | composite | 59 | -0.040 | 0.7225 |
| zReHo | axcpt | 53 | 0.045 | 0.4277 |
| zReHo | antisaccade | 48 | -0.302 | 0.0010 |
| zReHo | stopsignal | 77 | 0.096 | 0.2599 |
| zReHo | flanker | 91 | 0.058 | 0.4525 |
| zReHo | composite | 59 | 0.119 | 0.1254 |

**Table S11. All spatial correspondence tests with connectivity Gradients 1–3.**

| **Gradient** | **Metric** | **Task** | **n** | **ρ** | **Spin p** | **q (G1–G3)** |
| --- | --- | --- | --- | --- | --- | --- |
| G1 | BEN | age | 101 | 0.364 | 0.0002 | 0.0006 |
| G1 | BEN | axcpt | 53 | -0.117 | 0.2226 | 0.5567 |
| G1 | BEN | antisaccade | 48 | -0.046 | 0.5965 | 0.5965 |
| G1 | BEN | stopsignal | 77 | -0.250 | 0.0002 | 0.0006 |
| G1 | BEN | flanker | 91 | 0.048 | 0.6241 | 0.7013 |
| G1 | BEN | composite | 59 | -0.085 | 0.4561 | 0.4561 |
| G1 | ALFF | age | 101 | 0.103 | 0.5425 | 0.5425 |
| G1 | ALFF | axcpt | 53 | 0.004 | 0.9792 | 0.9792 |
| G1 | ALFF | antisaccade | 48 | 0.066 | 0.6107 | 0.7668 |
| G1 | ALFF | stopsignal | 77 | 0.097 | 0.5259 | 0.7888 |
| G1 | ALFF | flanker | 91 | -0.059 | 0.7283 | 0.7283 |
| G1 | ALFF | composite | 59 | -0.147 | 0.5103 | 0.5103 |
| G1 | fALFF | age | 101 | 0.100 | 0.3039 | 0.4562 |
| G1 | fALFF | axcpt | 53 | -0.123 | 0.4203 | 0.6305 |
| G1 | fALFF | antisaccade | 48 | -0.019 | 0.9018 | 0.9912 |
| G1 | fALFF | stopsignal | 77 | 0.028 | 0.7550 | 0.7550 |
| G1 | fALFF | flanker | 91 | -0.027 | 0.7894 | 0.9196 |
| G1 | fALFF | composite | 59 | -0.274 | 0.0812 | 0.1218 |
| G1 | ReHo | age | 101 | 0.007 | 0.9478 | 0.9478 |
| G1 | ReHo | axcpt | 53 | -0.084 | 0.3195 | 0.4543 |
| G1 | ReHo | antisaccade | 48 | -0.322 | 0.0010 | 0.0030 |
| G1 | ReHo | stopsignal | 77 | -0.088 | 0.3051 | 0.3995 |
| G1 | ReHo | flanker | 91 | 0.026 | 0.7672 | 0.7672 |
| G1 | ReHo | composite | 59 | -0.018 | 0.8286 | 0.8286 |
| G1 | zBEN | age | 101 | 0.121 | 0.1142 | 0.1713 |
| G1 | zBEN | axcpt | 53 | -0.047 | 0.5929 | 0.8448 |
| G1 | zBEN | antisaccade | 48 | -0.066 | 0.4077 | 0.6116 |
| G1 | zBEN | stopsignal | 77 | -0.111 | 0.1884 | 0.2825 |
| G1 | zBEN | flanker | 91 | 0.060 | 0.4723 | 0.7361 |
| G1 | zBEN | composite | 59 | 0.007 | 0.9312 | 0.9312 |
| G1 | zALFF | age | 101 | 0.046 | 0.7273 | 0.7273 |
| G1 | zALFF | axcpt | 53 | -0.283 | 0.0008 | 0.0024 |
| G1 | zALFF | antisaccade | 48 | -0.257 | 0.0040 | 0.0120 |
| G1 | zALFF | stopsignal | 77 | 0.058 | 0.7029 | 0.9944 |
| G1 | zALFF | flanker | 91 | 0.058 | 0.5853 | 0.5853 |
| G1 | zALFF | composite | 59 | 0.063 | 0.4961 | 0.8428 |
| G1 | zfALFF | age | 101 | 0.111 | 0.1858 | 0.4211 |
| G1 | zfALFF | axcpt | 53 | 0.014 | 0.8984 | 0.8984 |
| G1 | zfALFF | antisaccade | 48 | -0.037 | 0.7257 | 0.7257 |
| G1 | zfALFF | stopsignal | 77 | 0.223 | 0.0024 | 0.0072 |
| G1 | zfALFF | flanker | 91 | 0.011 | 0.9182 | 0.9182 |
| G1 | zfALFF | composite | 59 | -0.092 | 0.3747 | 0.3747 |
| G1 | zReHo | age | 101 | -0.038 | 0.6845 | 0.6845 |
| G1 | zReHo | axcpt | 53 | 0.026 | 0.6451 | 0.8604 |
| G1 | zReHo | antisaccade | 48 | -0.279 | 0.0012 | 0.0036 |
| G1 | zReHo | stopsignal | 77 | 0.107 | 0.2028 | 0.6083 |
| G1 | zReHo | flanker | 91 | 0.048 | 0.5393 | 0.5393 |
| G1 | zReHo | composite | 59 | 0.047 | 0.5115 | 0.7672 |
| G2 | BEN | age | 101 | 0.158 | 0.0504 | 0.0756 |
| G2 | BEN | axcpt | 53 | 0.063 | 0.5967 | 0.5967 |
| G2 | BEN | antisaccade | 48 | -0.060 | 0.5341 | 0.5965 |
| G2 | BEN | stopsignal | 77 | -0.031 | 0.7680 | 0.7680 |
| G2 | BEN | flanker | 91 | -0.173 | 0.0732 | 0.2196 |
| G2 | BEN | composite | 59 | -0.262 | 0.0248 | 0.0570 |
| G2 | ALFF | age | 101 | -0.180 | 0.3375 | 0.5425 |
| G2 | ALFF | axcpt | 53 | -0.064 | 0.6165 | 0.9247 |
| G2 | ALFF | antisaccade | 48 | -0.124 | 0.3961 | 0.7668 |
| G2 | ALFF | stopsignal | 77 | 0.262 | 0.1214 | 0.3641 |
| G2 | ALFF | flanker | 91 | 0.106 | 0.5729 | 0.7283 |
| G2 | ALFF | composite | 59 | 0.262 | 0.3635 | 0.5103 |
| G2 | fALFF | age | 101 | -0.082 | 0.5403 | 0.5403 |
| G2 | fALFF | axcpt | 53 | -0.353 | 0.0226 | 0.0678 |
| G2 | fALFF | antisaccade | 48 | 0.003 | 0.9912 | 0.9912 |
| G2 | fALFF | stopsignal | 77 | 0.129 | 0.2416 | 0.3623 |
| G2 | fALFF | flanker | 91 | 0.012 | 0.9196 | 0.9196 |
| G2 | fALFF | composite | 59 | -0.536 | 0.0016 | 0.0048 |
| G2 | ReHo | age | 101 | 0.033 | 0.8168 | 0.9478 |
| G2 | ReHo | axcpt | 53 | -0.096 | 0.2657 | 0.4543 |
| G2 | ReHo | antisaccade | 48 | -0.065 | 0.4531 | 0.4531 |
| G2 | ReHo | stopsignal | 77 | -0.134 | 0.1520 | 0.3995 |
| G2 | ReHo | flanker | 91 | -0.084 | 0.3167 | 0.7672 |
| G2 | ReHo | composite | 59 | -0.203 | 0.0016 | 0.0048 |
| G2 | zBEN | age | 101 | 0.103 | 0.0738 | 0.1713 |
| G2 | zBEN | axcpt | 53 | 0.078 | 0.4075 | 0.8448 |
| G2 | zBEN | antisaccade | 48 | 0.142 | 0.0730 | 0.2190 |
| G2 | zBEN | stopsignal | 77 | -0.017 | 0.8846 | 0.8846 |
| G2 | zBEN | flanker | 91 | 0.041 | 0.5351 | 0.7361 |
| G2 | zBEN | composite | 59 | -0.146 | 0.0274 | 0.0822 |
| G2 | zALFF | age | 101 | -0.099 | 0.4951 | 0.7273 |
| G2 | zALFF | axcpt | 53 | -0.133 | 0.1458 | 0.2187 |
| G2 | zALFF | antisaccade | 48 | 0.071 | 0.4181 | 0.6272 |
| G2 | zALFF | stopsignal | 77 | 0.023 | 0.8982 | 0.9944 |
| G2 | zALFF | flanker | 91 | 0.087 | 0.4743 | 0.5853 |
| G2 | zALFF | composite | 59 | 0.059 | 0.5619 | 0.8428 |
| G2 | zfALFF | age | 101 | 0.096 | 0.2807 | 0.4211 |
| G2 | zfALFF | axcpt | 53 | -0.057 | 0.6999 | 0.8984 |
| G2 | zfALFF | antisaccade | 48 | -0.077 | 0.4569 | 0.6854 |
| G2 | zfALFF | stopsignal | 77 | 0.142 | 0.0742 | 0.1113 |
| G2 | zfALFF | flanker | 91 | 0.181 | 0.0864 | 0.2591 |
| G2 | zfALFF | composite | 59 | -0.253 | 0.0172 | 0.0516 |
| G2 | zReHo | age | 101 | 0.046 | 0.6401 | 0.6845 |
| G2 | zReHo | axcpt | 53 | 0.038 | 0.5179 | 0.8604 |
| G2 | zReHo | antisaccade | 48 | 0.050 | 0.6119 | 0.6119 |
| G2 | zReHo | stopsignal | 77 | -0.074 | 0.4595 | 0.6893 |
| G2 | zReHo | flanker | 91 | -0.117 | 0.1346 | 0.4037 |
| G2 | zReHo | composite | 59 | -0.105 | 0.1948 | 0.5843 |
| G3 | BEN | age | 101 | 0.091 | 0.2390 | 0.2390 |
| G3 | BEN | axcpt | 53 | 0.066 | 0.3711 | 0.5567 |
| G3 | BEN | antisaccade | 48 | 0.059 | 0.4137 | 0.5965 |
| G3 | BEN | stopsignal | 77 | 0.088 | 0.1720 | 0.2579 |
| G3 | BEN | flanker | 91 | -0.028 | 0.7013 | 0.7013 |
| G3 | BEN | composite | 59 | 0.168 | 0.0380 | 0.0570 |
| G3 | ALFF | age | 101 | 0.077 | 0.3901 | 0.5425 |
| G3 | ALFF | axcpt | 53 | -0.054 | 0.5287 | 0.9247 |
| G3 | ALFF | antisaccade | 48 | -0.023 | 0.7668 | 0.7668 |
| G3 | ALFF | stopsignal | 77 | 0.018 | 0.8706 | 0.8706 |
| G3 | ALFF | flanker | 91 | 0.049 | 0.6351 | 0.7283 |
| G3 | ALFF | composite | 59 | -0.185 | 0.0394 | 0.1182 |
| G3 | fALFF | age | 101 | 0.069 | 0.3041 | 0.4562 |
| G3 | fALFF | axcpt | 53 | 0.017 | 0.8462 | 0.8462 |
| G3 | fALFF | antisaccade | 48 | -0.114 | 0.1918 | 0.5753 |
| G3 | fALFF | stopsignal | 77 | -0.140 | 0.0380 | 0.1140 |
| G3 | fALFF | flanker | 91 | -0.014 | 0.8654 | 0.9196 |
| G3 | fALFF | composite | 59 | -0.038 | 0.6899 | 0.6899 |
| G3 | ReHo | age | 101 | 0.173 | 0.0342 | 0.1026 |
| G3 | ReHo | axcpt | 53 | -0.047 | 0.4543 | 0.4543 |
| G3 | ReHo | antisaccade | 48 | -0.156 | 0.0240 | 0.0360 |
| G3 | ReHo | stopsignal | 77 | -0.062 | 0.3995 | 0.3995 |
| G3 | ReHo | flanker | 91 | 0.026 | 0.7131 | 0.7672 |
| G3 | ReHo | composite | 59 | -0.033 | 0.6459 | 0.8286 |
| G3 | zBEN | age | 101 | -0.028 | 0.6567 | 0.6567 |
| G3 | zBEN | axcpt | 53 | 0.013 | 0.8448 | 0.8448 |
| G3 | zBEN | antisaccade | 48 | -0.033 | 0.6301 | 0.6301 |
| G3 | zBEN | stopsignal | 77 | 0.132 | 0.0412 | 0.1236 |
| G3 | zBEN | flanker | 91 | 0.023 | 0.7361 | 0.7361 |
| G3 | zBEN | composite | 59 | 0.083 | 0.1944 | 0.2915 |
| G3 | zALFF | age | 101 | 0.138 | 0.0958 | 0.2873 |
| G3 | zALFF | axcpt | 53 | 0.021 | 0.7748 | 0.7748 |
| G3 | zALFF | antisaccade | 48 | -0.034 | 0.6529 | 0.6529 |
| G3 | zALFF | stopsignal | 77 | 0.001 | 0.9944 | 0.9944 |
| G3 | zALFF | flanker | 91 | 0.073 | 0.3927 | 0.5853 |
| G3 | zALFF | composite | 59 | 0.010 | 0.8920 | 0.8920 |
| G3 | zfALFF | age | 101 | -0.004 | 0.9494 | 0.9494 |
| G3 | zfALFF | axcpt | 53 | 0.083 | 0.2551 | 0.7654 |
| G3 | zfALFF | antisaccade | 48 | -0.075 | 0.3449 | 0.6854 |
| G3 | zfALFF | stopsignal | 77 | -0.062 | 0.3291 | 0.3291 |
| G3 | zfALFF | flanker | 91 | 0.022 | 0.7708 | 0.9182 |
| G3 | zfALFF | composite | 59 | -0.072 | 0.3427 | 0.3747 |
| G3 | zReHo | age | 101 | 0.097 | 0.1674 | 0.5021 |
| G3 | zReHo | axcpt | 53 | 0.011 | 0.8604 | 0.8604 |
| G3 | zReHo | antisaccade | 48 | -0.124 | 0.0700 | 0.1050 |
| G3 | zReHo | stopsignal | 77 | -0.020 | 0.7752 | 0.7752 |
| G3 | zReHo | flanker | 91 | -0.075 | 0.2789 | 0.4184 |
| G3 | zReHo | composite | 59 | -0.007 | 0.9104 | 0.9104 |

### Supplementary figures

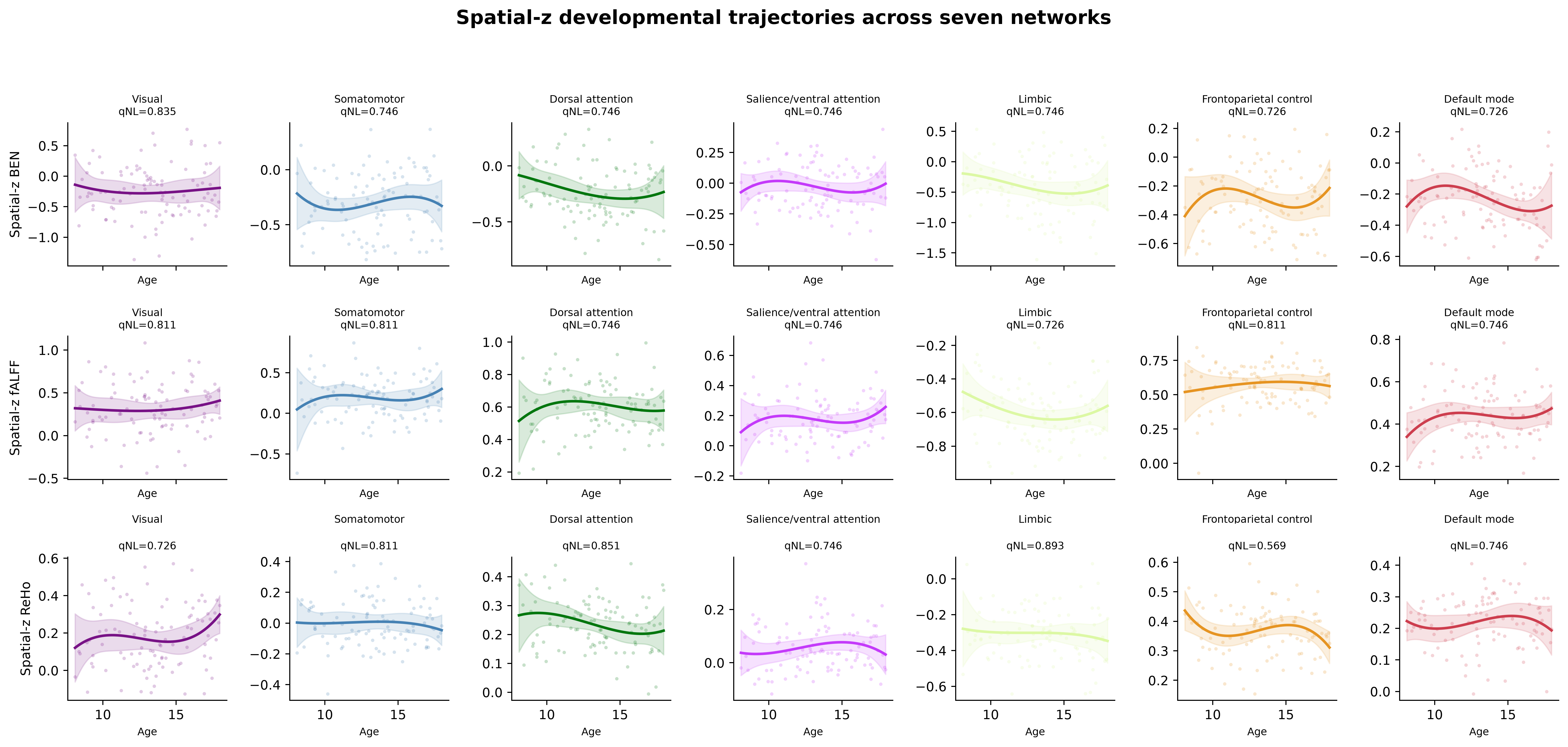

Figure S2. Seven-network developmental trajectories for spatial-z BEN, fALFF, and ReHo. Curves are cubic fits with HC3 95% confidence bands; FDR-significant omnibus age associations are reported in Table S7; curvature tests are shown in Figure S3.

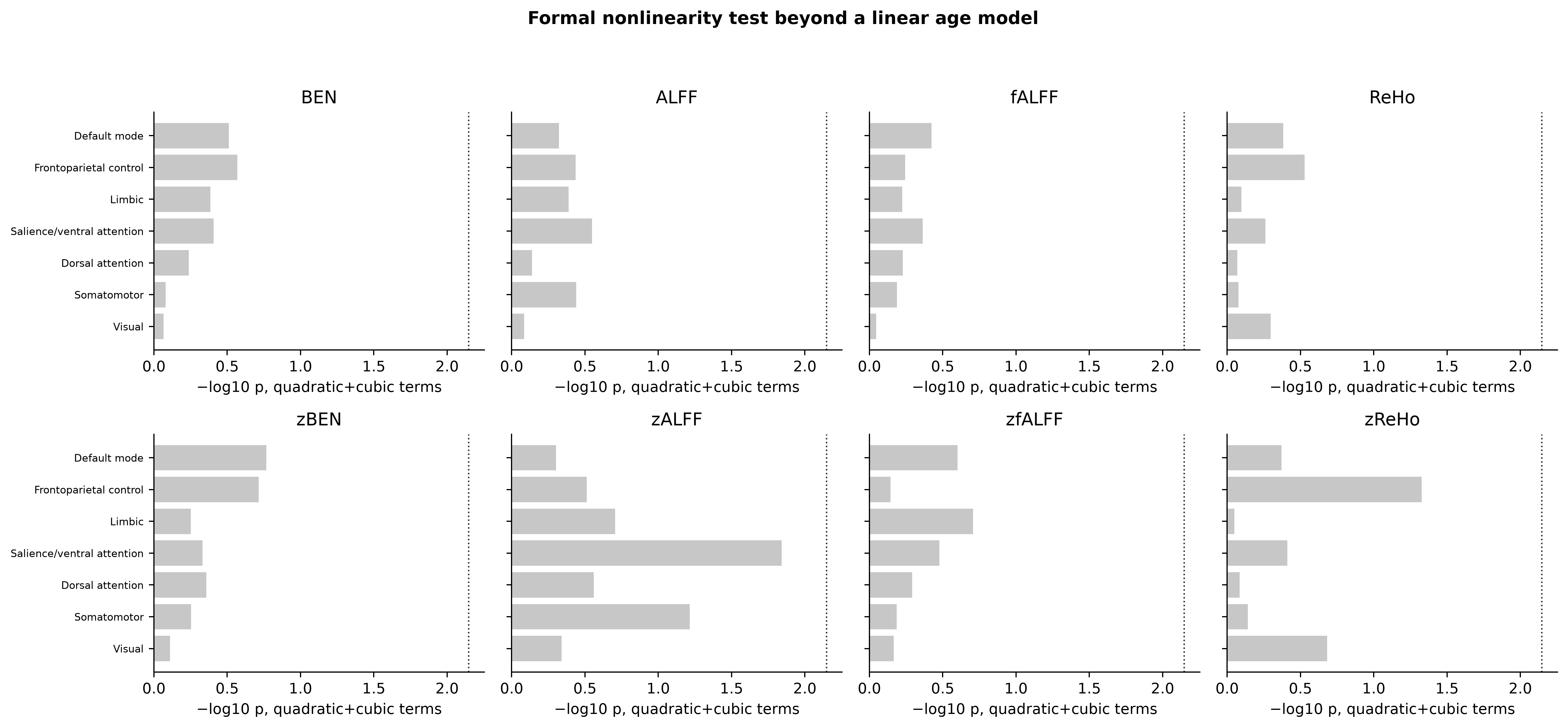

Figure S3. Tests of curvature beyond linear age across metrics and networks. No metric-network result survived seven-network FDR.

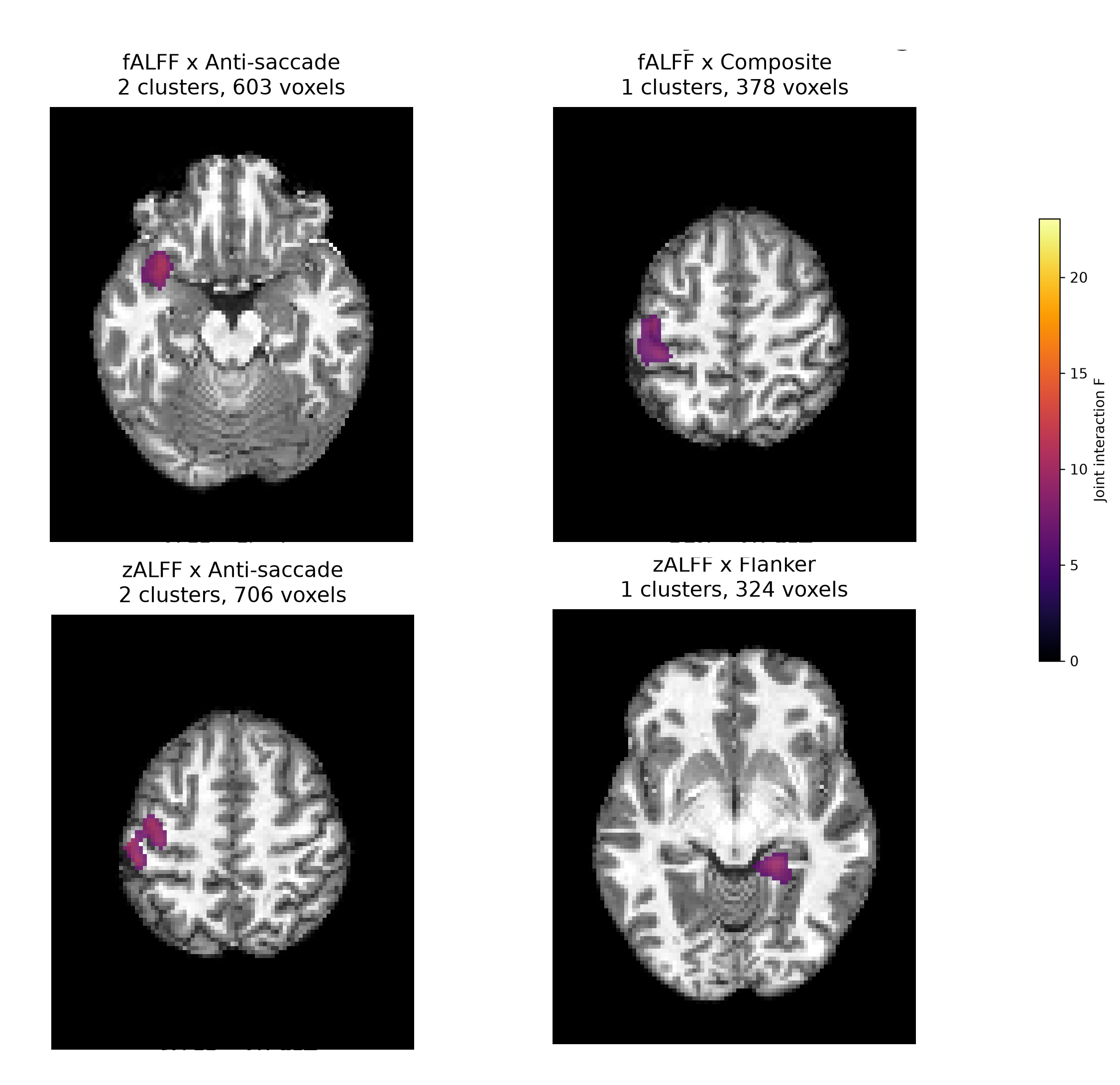

Figure S4. Corrected voxelwise task-by-age interactions for raw fALFF with anti-saccade and composite performance and spatial-z ALFF with anti-saccade and Flanker performance. Maps show joint interaction F statistics within clusters surviving permutation cluster-mass FDR in each metric-by-task family. AX-CPT maps are presented in main Figure 2.

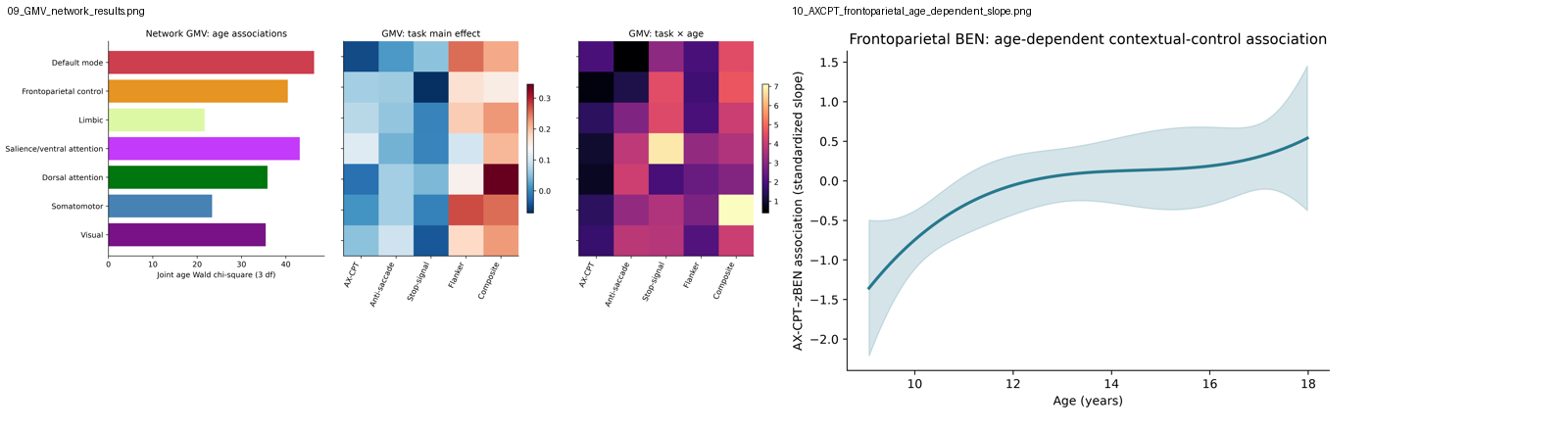

Figure S5. Network GMV analyses and the conditional age-varying frontoparietal BEN–AX-CPT slope.
